# Deficiency in cytosine DNA methylation leads to high chaperonin expression and tolerance to aminoglycosides in *Vibrio cholerae*

**DOI:** 10.1101/2021.07.29.454301

**Authors:** André Carvalho, Didier Mazel, Zeynep Baharoglu

## Abstract

Antibiotic resistance has become a major global issue. Understanding the molecular mechanisms underlying microbial adaptation to antibiotics is of keen importance to fight Antimicrobial Resistance (AMR). Aminoglycosides are a class of antibiotics that target the small subunit of the bacterial ribosome, disrupting translational fidelity and increasing the levels of misfolded proteins in the cell. In this work, we investigated the role of VchM, a DNA methyltransferase, in the response of the human pathogen *Vibrio cholerae* to aminoglycosides. VchM is a *V. cholerae* specific orphan m5C DNA methyltransferase that generates cytosine methylation at 5′-R**<u>C</u>**CGGY-3′ motifs. We show that deletion of *vchM*, although causing a growth defect in absence of stress, allows *V. cholerae* cells to cope with aminoglycoside stress at both sub-lethal and lethal concentrations of these antibiotics. Through transcriptomic and genetic approaches, we show that *groESL-2* (a specific set of chaperonin-encoding genes located on the second chromosome of *V. cholerae*), are upregulated in cells lacking *vchM* and are needed for the tolerance of *vchM* mutant to lethal aminoglycoside treatment, likely by fighting aminoglycoside-induced misfolded proteins. Interestingly, preventing VchM methylation of the four RCCGGY sites located in *groESL-2* region, leads to a higher expression of these genes in WT cells, showing that VchM modulates the expression of these chaperonins in *V. cholerae* directly through DNA methylation.

**AUTHOR SUMMARY:** Bacteria are organisms with a remarkable ability to adapt to several stress conditions, including to the presence of antibiotics. The molecular mechanisms underlying such adaptation lead, very often, to phenomena like antimicrobial tolerance and resistance, responsible for the frequent failure of antibiotic treatment. The study of these molecular mechanisms is thus an important tool to understand development of antimicrobial resistance in bacteria. In this work, we show that abrogating cytosine DNA methylation in *Vibrio cholerae* increases its tolerance to aminoglycosides, a class of antibiotics that cause protein misfolding. DNA methylation is known to affect gene expression and regulate several cellular processes in bacteria. Here we provide evidence that DNA methylation also has a more direct role in controlling antibiotic susceptibility in bacteria. Consequently, the study of bacterial DNA methyltransferases and DNA methylation should not be overlooked when addressing the problem of antimicrobial tolerance/resistance.

## INTRODUCTION

In the past decades, the over/misuse and large-scale production of antibiotics has created a serious ecological problem with important consequences for the emergence of antimicrobial resistance (AMR). In fact, a large proportion of the antibiotics ingested are released intact in the environment (1, 2) and found at trace levels or as gradients in various environments (3, 4). Hence, in these environments, one can find the presence of very low doses of drugs commonly referred as subMIC, i.e. under the MIC (Minimal Inhibitory Concentration). Although not enough to kill or prevent the growth of bacterial populations, subMIC doses of antibiotics are proposed to work as signaling molecules (5) and trigger important stress mechanisms that often result in development of antibiotic resistance (4, 6–8). We have previously shown that subMIC of antibiotics, such as aminoglycosides, trigger common and specific stress responses in Gram-negative bacteria (9, 10).

Aminoglycosides (AGs) are positively charged molecules that bind 16S rRNA at the 30S ribosomal subunit and negatively affect translation. Specifically, AGs (e.g. tobramycin, streptomycin, kanamycin, gentamicin and neomycin) are known to disrupt translational fidelity and increase the levels of mistranslation, i.e. the misincorporation of certain amino acids in proteins (11, 12). In turn, high levels of mistranslation result in the production and accumulation of aberrant proteins in the cell, which contribute to the collapse of important cell processes and ultimately lead to cell death (13, 14).

*V. cholerae* is a water-borne gram-negative bacterium, human pathogen and the causative agent of cholera disease. As part of its life cycle, *V. cholerae* often transits between the human gut and the external environment where it can find low doses of antibiotics. During our studies to better understand adaption of *V. cholerae* to aminoglycosides (15), we observed that a mutant of a *V*. cholerae’s specific DNA methyltransferase (*vca0198* - VchM) was less susceptible to aminoglycosides than its isogenic WT strain, suggesting that DNA methylation could play a role in *V. cholerae* adaptation to AGs. *vchM* codes for an Orphan m5C DNA methyltransferase that causes DNA methylation at 5′-RCCGGY-3′ motifs (16). DNA methylation is catalyzed by enzymes called DNA methyltransferases (DNA MTases) that transfer a methyl group from S-adenosyl-L methionine (SAM) to adenine and cytosine in specific DNA motifs (17, 18). As a result, one can find the existence of small amounts of N6-methyl-adenine (6mA), C5-methyl-cytosine (5mC) and N4-methyl-cytosine (4mC) in the DNA of both eukaryotes and prokaryotes. In bacteria, the existence of such modified DNA bases have been shown to play a critical role in processes such as protection against invasive DNA, DNA replication and repair, cell cycle regulation and control of gene expression (19–23).

While it was previously proposed that VchM plays a role in the cell envelope stress response of *V. cholerae* (23), no link between this DNA MTase and antibiotic stress has yet been established. Here, we show that deletion of *vchM* (although causing a growth defect in absence of stress) allows *V. cholerae* cells to better deal with the effect of aminoglycosides. In fact, not only the *vchM* mutant is a better competitor during growth in presence of subMIC doses of aminoglycosides, it is also more tolerant to killing by lethal doses of these antibiotics. Transcriptome analysis of a Δ*vchM* strain revealed the upregulation of *groESL-2* genes, a specific set of chaperonin-encoding genes located on the second chromosome of *V. cholerae*. High expression of *groESL-2* genes (but not of chromosome one *groESL-1* homologues) determines the higher tolerance of Δ*vchM* to lethal AG treatment, suggesting a new and specific role of *groESL-2* in managing AG-mediated proteotoxic stress. Interestingly, we observed the presence of four VchM motifs in *groESL-2* region. Preventing methylation of all these sites in the WT strain by disrupting such motifs results in increased expression of these genes. Intriguingly, the high expression of *groESL-2* does not seem to contribute to the competitive advantage of the Δ*vchM* strain grown under subMIC AG which suggests the involvement of additional players in the global response of Δ*vchM* to aminoglycosides.

## RESULTS

### *V. cholerae* cells lacking *vchM* cope better with subMIC doses of AGs

In order to explore a possible role of *vchM* in the response of *V. cholerae* O1 El Tor N16961 to aminoglycosides, we constructed an in-frame deletion mutant of *vchM* by allelic replacement with an antibiotic resistance cassette, and compared its growth to the isogenic wild-type (WT) strain, in rich media, with or without increasing concentrations of subMIC tobramycin (Fig 1). As previously described (23), this mutant exhibits a reduced doubling rate when grown in monoculture in antibiotic free rich media. However, the difference in growth between WT and Δ*vchM* strains observed in absence of antibiotics becomes gradually more negligible with increasing concentrations of subMIC TOB. At higher concentrations (90% of the MIC), Δ*vchM* even displays a clear advantage over the WT (Fig 1A). Importantly, a Δ*vchM* strain harboring a low-copy number plasmid with *vchM* gene under the control of its own promoter behaves as the WT strain in absence of tobramycin and even slightly worse in presence of higher doses of this drug (Fig 1A), showing that the observed growth phenotypes are due to the absence of *vchM*.

**Fig 1.**
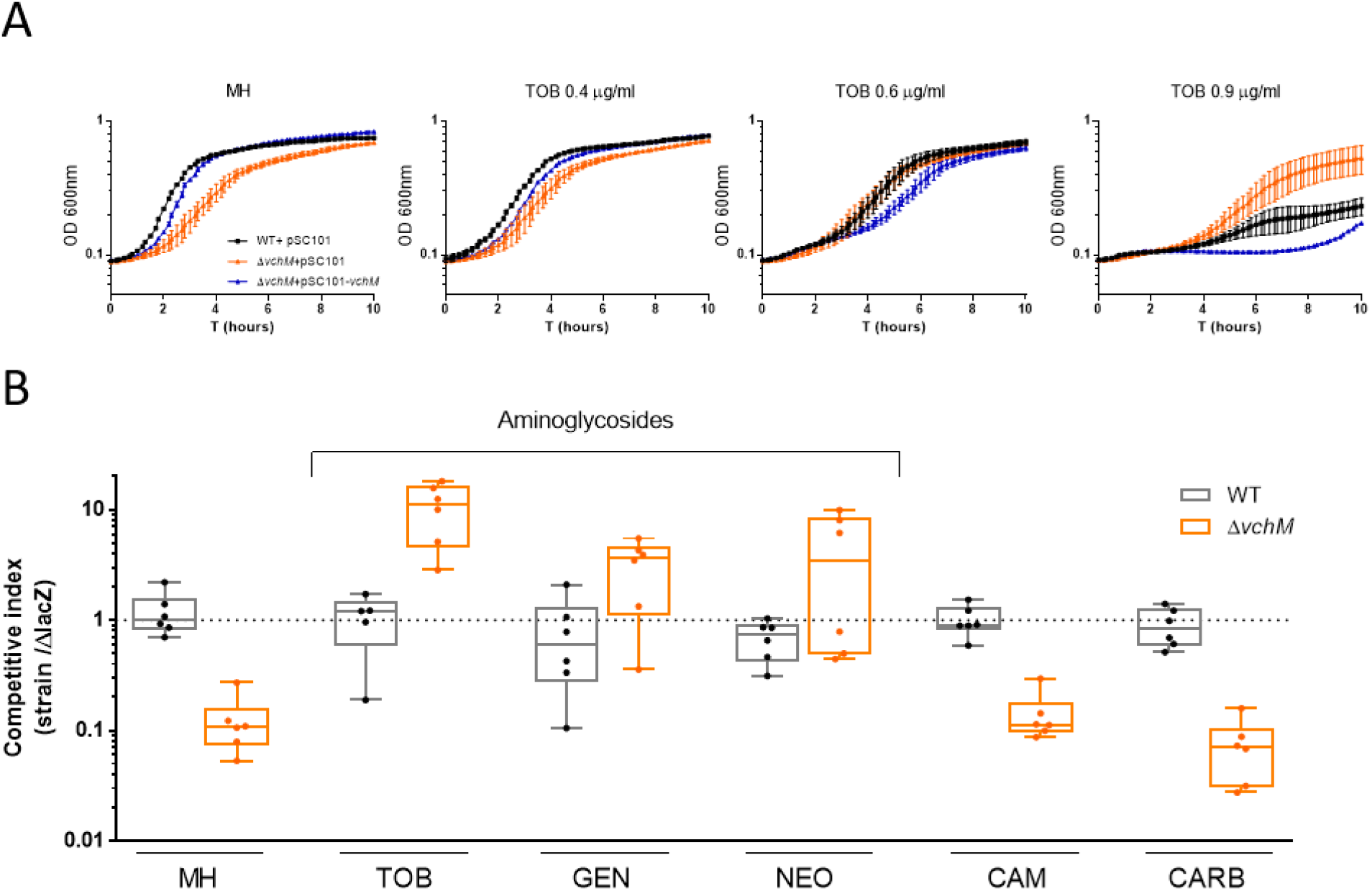
*V. cholerae* N16961 Δ*vchM* is less susceptible to subMIC aminoglycosides. **A**. Growth curves in absence (MH) or presence of subMIC doses of tobramycin. Bars represent SD (n=3) **B.** *In vitro* competitions of WT and mutant strains against isogenic Δ*lacZ* reference strain in absence or presence of different antibiotics at subMIC concentrations (TOB, 0.6 μg/ml; GEN, 0.5 μg/ml; NEO, 2.0 μg/ml; CAM, 0.4 μg/ml; CARB, 2.5 μg/ml). Box plots indicate the median and the 25^th^ and 75^th^ percentiles; whiskers indicate the min and max values (n=6).

Next, we asked whether the growth phenotype observed in monocultures was translatable to a higher relative fitness in co-cultures in the presence of subMIC doses of tobramycin and other AGs. For that, we competed both WT and Δ*vchM* strains (both *lacZ^+^*) with an isogenic Δ*lacZ* mutant (initial ratio of 1:1), in MH or MH supplemented with subMIC concentrations (50% MIC) of the aminoglycosides tobramycin (TOB), gentamicin (GEN) and neomycin (NEO). We assessed relative fitness by plating cultures after 20 hours of growth and counting the final proportion of lacZ^+^/lacZ^-^ colonies. Competition of WT against the *lacZ^-^* mutant served as a control to account for any effect of *lacZ* deletion on growth. Supporting the previous results in monocultures, Δ*vchM* is outcompeted by the *lacZ* mutant in MH (≈10-fold difference) (Fig 1B). More importantly, in presence of low concentrations of aminoglycosides, Δ*vchM* is either equally competitive or even displays a clear growth advantage over the reference strain (Fig 1B). Additionally, in order to test whether these results hold for drugs other than aminoglycosides we performed competitions in the presence of chloramphenicol (CAM) and the beta-lactam carbenicillin (CARB). Unlike AGs, the presence of low concentrations of these drugs did not increased the relative fitness of the Δ*vchM* mutant.

Altogether, these results confirm that lack of *vchM* in *V. cholerae* negatively impacts growth in antibiotic-free media (23) but confers a selective advantage to *V. cholerae* in presence of subMIC doses of AGs (Fig 1). In order to test if deleting *vchM* affects the MIC of these drugs, we measured the MIC of both WT and Δ*vchM* mutant and found no difference (Table 1).

**Table 1.** MICs (μg/ml) of the different antibiotics tested for *V. cholerae* N16961 WT and *ΔvchM* strains

| Strain | TOB | GEN | NEO | CAM | CARB |
| --- | --- | --- | --- | --- | --- |
| WT | 1-2 | 1 | 4 | 1 | 15 |
| *ΔvchM* | 1-2 | 1 | 4 | 1 | 7.5 |

TOB, tobramycin; GEN, gentamicin; NEO, neomycin; CAM, chloramphenicol; CARB, carbenicillin

### VchM deficiency promotes higher tolerance to lethal doses of aminoglycosides

It has been previously shown that certain mutations affect functions conferring bacterial populations a tolerant phenotype towards a specific drug (24). Such bacterial populations can transiently withstand lethal doses of that drug without necessarily any impact on the MIC of the population (24).

To continue exploring the different susceptibility of the Δ*vchM* strain to aminoglycosides we assessed the survival rate of *V. cholerae* WT and Δ*vchM* strains during treatment with lethal doses of tobramycin and gentamicin at 20x and 10x the MIC, respectively (Fig 2). Given the inherent growth defect of VchM deficiency, we performed time-dependent killing curves on stationary phase cells, where both WT and Δ*vchM* strains are no longer actively growing, excluding a possible link between growth rate and aminoglycoside lethality as previously shown (25). Strikingly, survival to both antibiotics was increased 10-1000 fold in the Δ*vchM* mutant, suggesting that the absence of VchM allows *V. cholerae* to transiently withstand lethal doses of these aminoglycosides (Fig 2).

**Fig 2.**
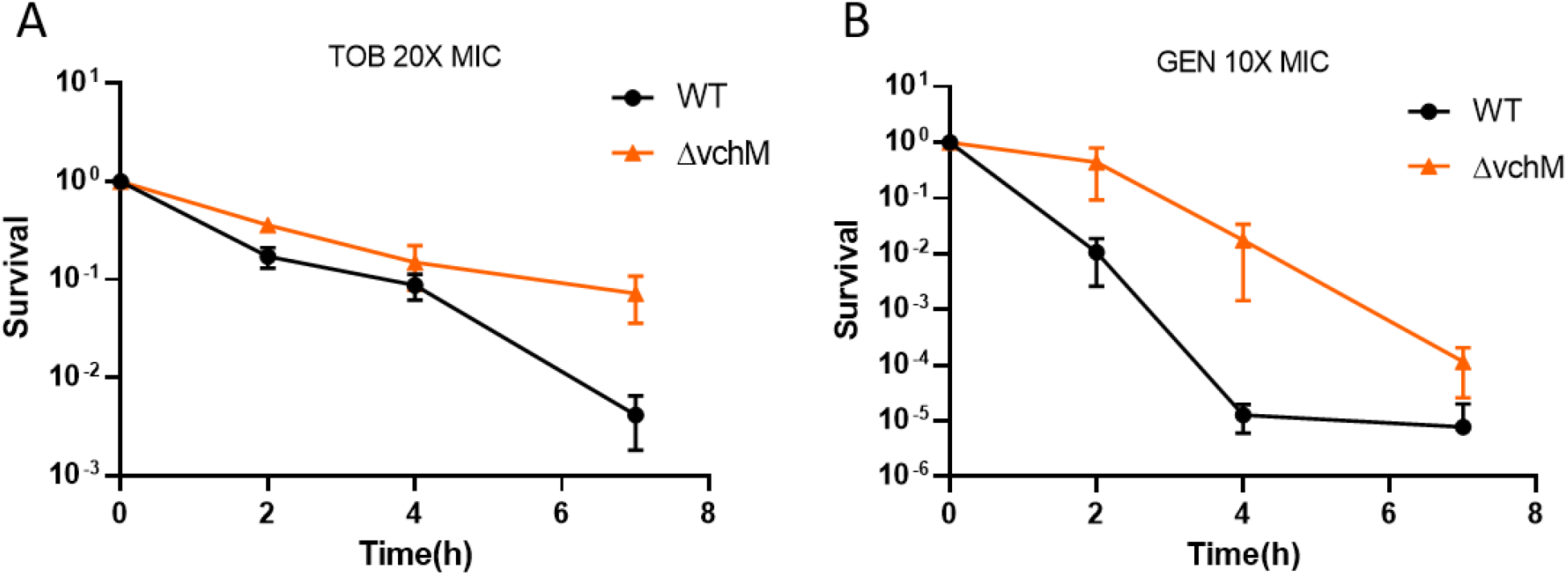
Δ*vchM* strain is more tolerant to lethal aminoglycoside treatment. Survival of stationary-phase WT and Δ*vchM* cells exposed to lethal doses of tobramycin (TOB) **(A)**, and gentamicin (GEN) **(B)**. Survival represents the number of bacteria (CFU/mL) after treatment divided by the initial number of bacteria prior treatment. Means and SD are represented, n=3.

One crucial aspect that determines the efficacy of aminoglycoside treatment is the uptake of these drugs by the bacterial cell. This process is energy dependent and requires a threshold membrane potential (26). We used a previously reported assay that measures cell fluorescence after incubation with fluorescent-marked neomycin (neo-cy5) (27) as a proxy for aminoglycoside uptake. We did not observe any difference in fluorescence between WT and Δ*vchM* mutant strains (S1 Fig). Thus, differential uptake of aminoglycosides is unlikely the reason for the increased tolerance to these drugs in Δ*vchM*.

### A specific set of chaperonins is upregulated in Δ*vchM* cells

To understand the high tolerance to aminoglycosides observed in Δ*vchM*, we performed RNA-seq on stationary phase cells of WT and Δ*vchM* strains grown in rich, stress-free media. The analysis of the transcriptome of Δ*vchM V. cholerae* O1 El Tor N16961 strain reveals the significant upregulation (fold change ≥ 2, p<0.01) and downregulation (fold change ≤ −2, p<0.01) of 68 and 53 genes, respectively (S1 Table). Among the differentially expressed, we found four genes directly involved in protein folding to be upregulated in Δ*vchM* strain. Those are the molecular chaperones GroEL and co-chaperonins GroES (Table 2). In many bacterial species, GroEL and its co-chaperonin GroES form a molecular machine essential for folding of large newly synthetized proteins also helping re-folding of proteins damaged by proteotoxic stress (28). Interestingly, overexpression of GroES and GroEL proteins was found to promote short-term tolerance to aminoglycoside-induced protein misfolding in *E. coli* (29).

**Table 2.** Protein folding and stabilization genes upregulated in Δ*vchM* (fold change > 2, p-value < 0.01)

| Locus | Name | Fold change<br>($\Delta vchM$/WT) | Annotation |
| --- | --- | --- | --- |
| vc2665 | groEL-1 | **2.24** | Chaperonin, 60-kDa subunit |
| vca0820 | groEL-2 | **3.54** | Chaperonin, 60-kDa subunit |
| vc2664 | groES -1 | **2.60** | Chaperonin, 10-kDa subunit |
| vca0819 | groES-2 | **6.78** | Chaperonin, 10-kDa subunit |

*V. cholerae* is one of, at least, seven *Vibrio* species harboring two copies of *groES – groEL* (*groESL*) bicistronic operons (30). Whereas *groESL-1* is encoded in chromosome 1 (*vc2664-vc2665*), *groESL-2* is located in chromosome 2 (*vca0819-0820*) (Fig 3A) (30). Based on our RNA-seq data, the latter manifested a larger fold change (Table 2). In order to confirm differential expression of these genes in Δ*vchM*, we measured *groES-1* and *groES-2* relative gene expression in exponential and stationary phase cells of WT and mutant strains, using digital qRT-PCR with the housekeeping *gyrA* gene as reference (31). The results confirm a higher induction of *groES-2* genes in both exponential and stationary phase Δ*vchM* cells with a fold change (over the WT) of ca. 10X and 5X, respectively (Fig 3B). However, *groES-1* fold change was only slightly increased in exponential and unnoticeable in stationary phase.

**Fig 3.**
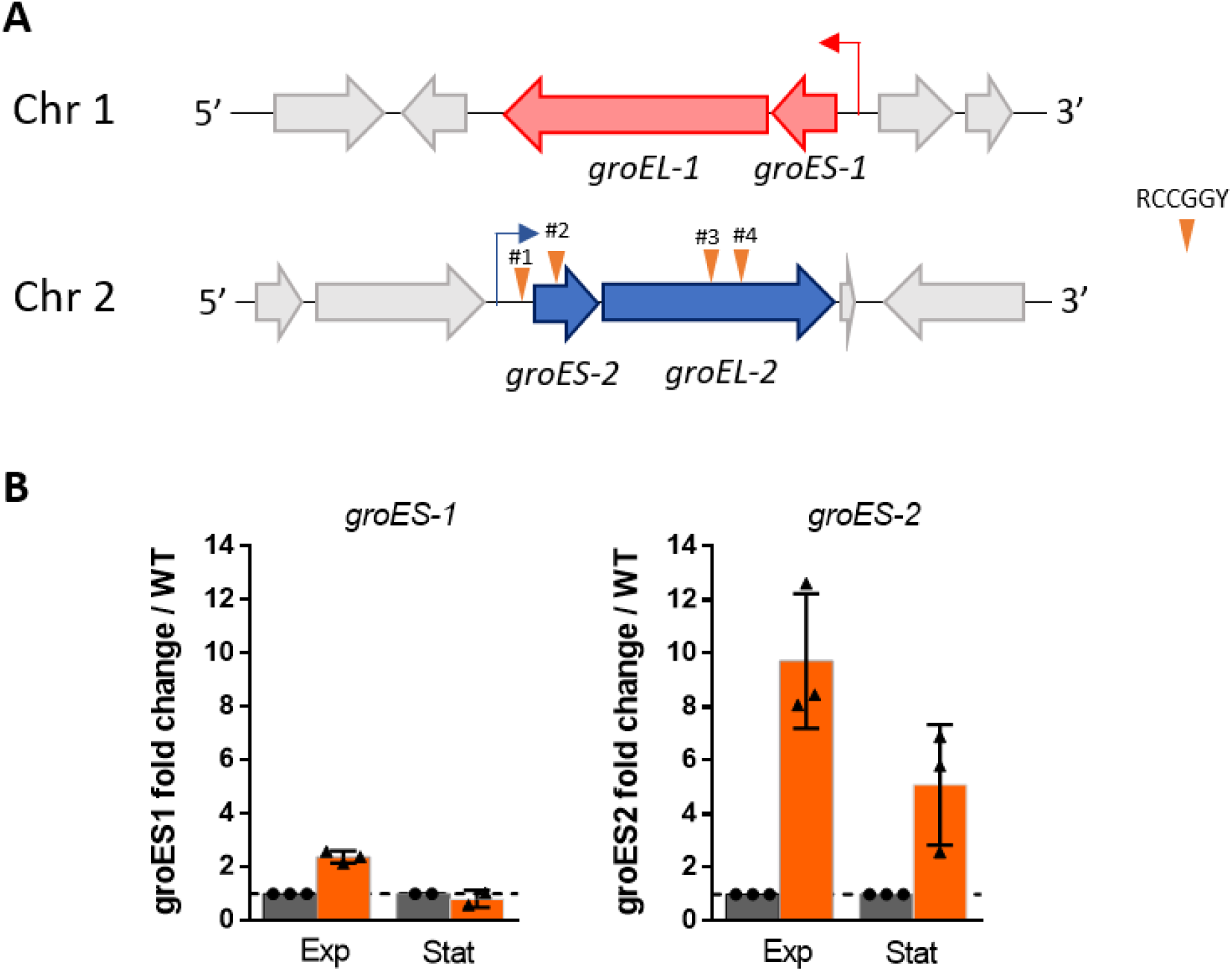
*groESL-2* operon is upregulated in Δ*vchM* strain. **A.** Schematic representation of both *groESL* operons in *V. cholerae*. The four RCCGGY sites present along the *groESL-2* region are represented by the inverted orange triangles. **B.** Fold change (Δ*vchM*/WT) of the relative expression levels of *groES-1* and *groES-2* in cultures at exponential phase (Exp, OD_600_ ≈ 0.3) or stationary phase (Stat, OD_600_ ≈ 1.8-2.0). Means and SD are represented, n=3.

Induction of groESL genes is usually associated to perturbations in proteostasis which leads to activation of the heat-shock response (32). Indeed, expression of both *groESL-1* and *groESL-2* is controlled by the heat-shock alternative sigma factor RpoH in *V. cholerae* (33). However, the upregulation of *groESL-2* genes in Δ*vchM* cells is likely independent of heat-shock activation as i) we do not observe any other genes of the heat-shock regulon being upregulated in the mutant (for example, *dnaKJ/grpE, clpB, ibpAB*) and ii) expression of *groESL-1* is not as increased as expression of *groESL-2* (Fig 3B).

Altogether, these results confirm that in absence of VchM, expression of *groESL-2* genes is markedly increased in *V. cholerae* and suggest that regulation of *groESL-2* operon in the Δ*vchM* mutant can be independent of heat-shock response.

### Deletion of *groESL-2* operon abolishes Δ*vchM* high tolerance to lethal doses of tobramycin

In bacteria that harbor a single copy of this operon, GroESL are essential proteins for cell viability (34). However, possible redundancy between *groESL-1* and *groESL-2* could allow for the deletion of one or the other operon in *V. cholerae*. Thus, we attempted to delete *groESL-1* and *groESL-2* from *V. cholerae* WT and Δ*vchM* strains. While Δ*groESL-2* and Δ*vchM groESL-2* strains were easily obtained, we could not manage to delete *groESL-1* in any background despite several attempts. Moreover, deletion of *groESL-2* did not affect the growth of V*. cholerae* in rich medium (Fig 4A). Respectively, GroES-1 and GroEL-1 share 80% and 87% amino acid identity with the only and essential GroES and GroEL proteins of *E. coli*, but lower (66% and 76%) amino acid identity with GroES-2 and GroEL-2 (S2 Fig). These observations suggest that i) GroESL-1 (but not GroESL-2) is essential for *V. cholerae* viability and ii) GroESL-1 is probably the main housekeeping chaperonin system while the divergent GroESL-2 could act synergistically in response to high levels of misfolding or having specific substrates upon protein damage caused by specific stresses. Surprisingly, competition of Δ*groESL-2* with a lacZ^-^ strain shows that loss of these proteins is not detrimental for growth of *V. cholerae* in presence of subMIC TOB (S3A Fig). Similarly, survival of the Δ*groESL-2* strain to lethal doses of TOB does not differ from that of the WT (S3B Fig). However, these genes are intrinsically highly expressed in Δ*vchM* strain, where they may confer a selective advantage in presence of AG stress. In this case, deletion of *groESL-2* in Δ*vchM* background would affect the mutant’s tolerance. Indeed, when we compared the survival to lethal AG treatment of Δ*vchM* to that of a Δ*vchM groESL-2* double mutant, we found that the absence of *groESL-2* abolishes high tolerance to tobramycin and gentamicin in Δ*vchM* (Fig 4B), without affecting growth in absence of stress (Fig 4A). These results show that the higher expression of *groESL-2* is required for the high tolerance of the Δ*vchM* mutant to lethal AG treatment.

**Fig 4.**
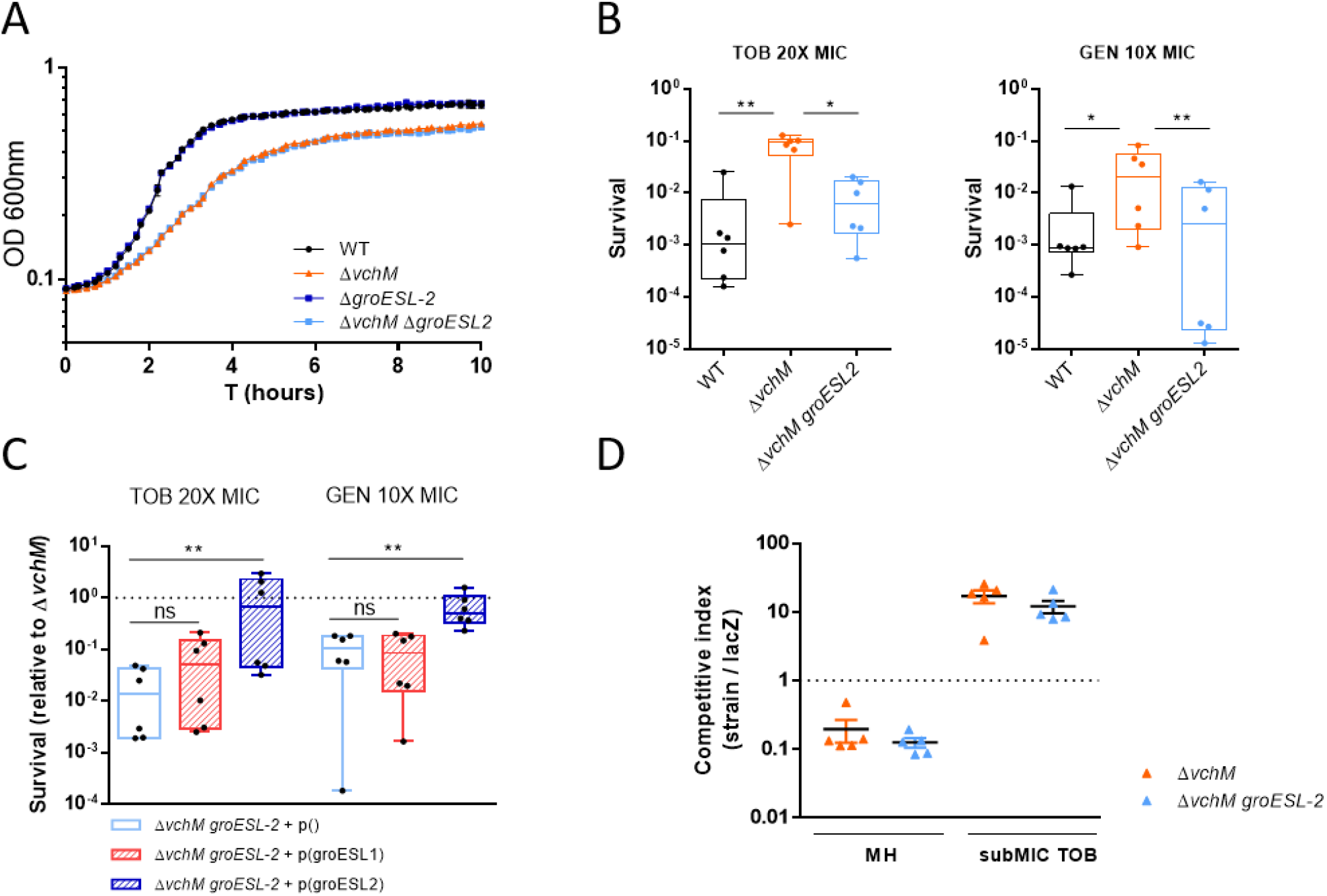
*groESL-2* is needed for the increased tolerance of Δ*vchM* to lethal AG treatment. **A.** Growth curves in MH medium. Means and SD are represented, n=3. **B.** Survival of stationary-phase WT and Δ*groESL-2* cells exposed to lethal aminoglycoside treatment for 7 hours. Box plots indicate the median and the 25^th^ and 75^th^ percentiles; whiskers indicate the min and max values (n=6 from two independent experiments). **C.** Survival (after 7 hours AG treatment) of Δ*vchM groESL-2* double mutant harboring an empty plasmid or a plasmid expressing either *groESL-1* or *groESL-2*, relative to survival of the Δ*vchM* with the control plasmid. Box plots indicate the median and the 25^th^ and 75^th^ percentiles; whiskers indicate the min and max values (n=6 from two independent experiments). In **B** and **C** statistically significant differences were determined using Friedman’s test with Dunn’s post-hoc test for multiple comparisons. * P<0.05, ** P<0.01, ns = not significant. **D.** *In vitro* competitions of Δ*vchM* and Δ*vchM groESL-2* double mutant strains against isogenic Δ*lacZ* reference strain in absence or presence of subMIC TOB, 0.6 μg/ml; Error bars indicate SD (n=6).

We then tested whether the high tolerance of this mutant relies on general higher levels of chaperonins or if it specifically linked to GroESL-2 chaperonins. We thus tried to complement the Δ*vchM groESL-2* mutant by ectopically expressing *groESL-1* or *groESL-2* and assessed survival to lethal doses of AGs. Strikingly, only overexpression of *groESL-2* is able to promote survival levels similar to those observed in Δ*vchM* (Fig 4C), suggesting a specific role for GroESL-2 in managing AG-mediated proteotoxic stress in *V. cholerae* cells lacking VchM. Interestingly, we observed no difference in the relative fitness of Δ*vchM* and Δ*vchM groESL-2* mutants in competitions in presence of subMIC doses of tobramycin, which shows that *groESL-2* it is not implicated in Δ*vchM* higher relative fitness during growth in subMIC AGs (Fig 4D).

### VchM controls *groESL-2* expression through direct DNA methylation

Knowing the role of VchM in regulating gene expression in *V. cholerae* (23), we asked whether VchM controls *groESL* expression directly through DNA methylation. VchM methylates the first cytosine in 5′-R**C**CGGY-3′ motifs (16). This prompted us to search for such motifs in both *groESL* operons. While we couldn’t detect any of these sites along the *groESL-1* locus, we found a total of four VchM motifs in *groESL-2* region: motif #1 within the 5′ UTR of the operon, 47 bp away from the initiation codon; motif #2 is within the coding region of *groES-2* while motifs #3 and #4 are located within the coding region of *groEL-2* (Fig 5A). We hypothesized that the methylation state of these motifs could modulate the transcription of *groESL-2* genes. To test this, we generated a mutant by replacing all RCCGGY motifs in *groESL-2* region by non-consensus motifs but maintaining the amino acid sequence of GroESL-2 proteins intact (Fig 5A, mut#1-4). Additionally, we created a mutant where only the RCCGGY #1 was altered in order to investigate if this site, for being in the regulatory region of this operon, had a stronger contribution in modulating gene expression (Fig 5A, mut#1). We then measured *groES-2* expression in both mutants and observed that disruption of RCCGGY #1 lead to a very weak increase in *groES-2* expression relative to the WT, while disruption of all four sites led to a significantly higher expression of this gene (Fig 5B). We additionally tested *groEL-2* expression and observed similar results (S4 Fig). Supporting our hypothesis that this regulation is methylation-dependent, we did not observe any difference in *groES-2* or *groEL-2* expression when we mutated sites #1-4 in the Δ*vchM* background (S4 Fig). It is worth mentioning that, in these experiments, the expression of *groES-2* in the Δ*vchM* strain was consistently higher than in the WT mut#1-4 (Fig 5B) suggesting that an additional factor, in synergy with the methylation of RCCGGY sites, may control expression of *groES-2*. Nonetheless, overall these results show that a specific set of chaperonin encoding genes is under the control of DNA cytosine methylation in *V. cholerae*, linking DNA methylation to modulation of chaperonin expression and tolerance to antibiotics.

**Fig 5.**
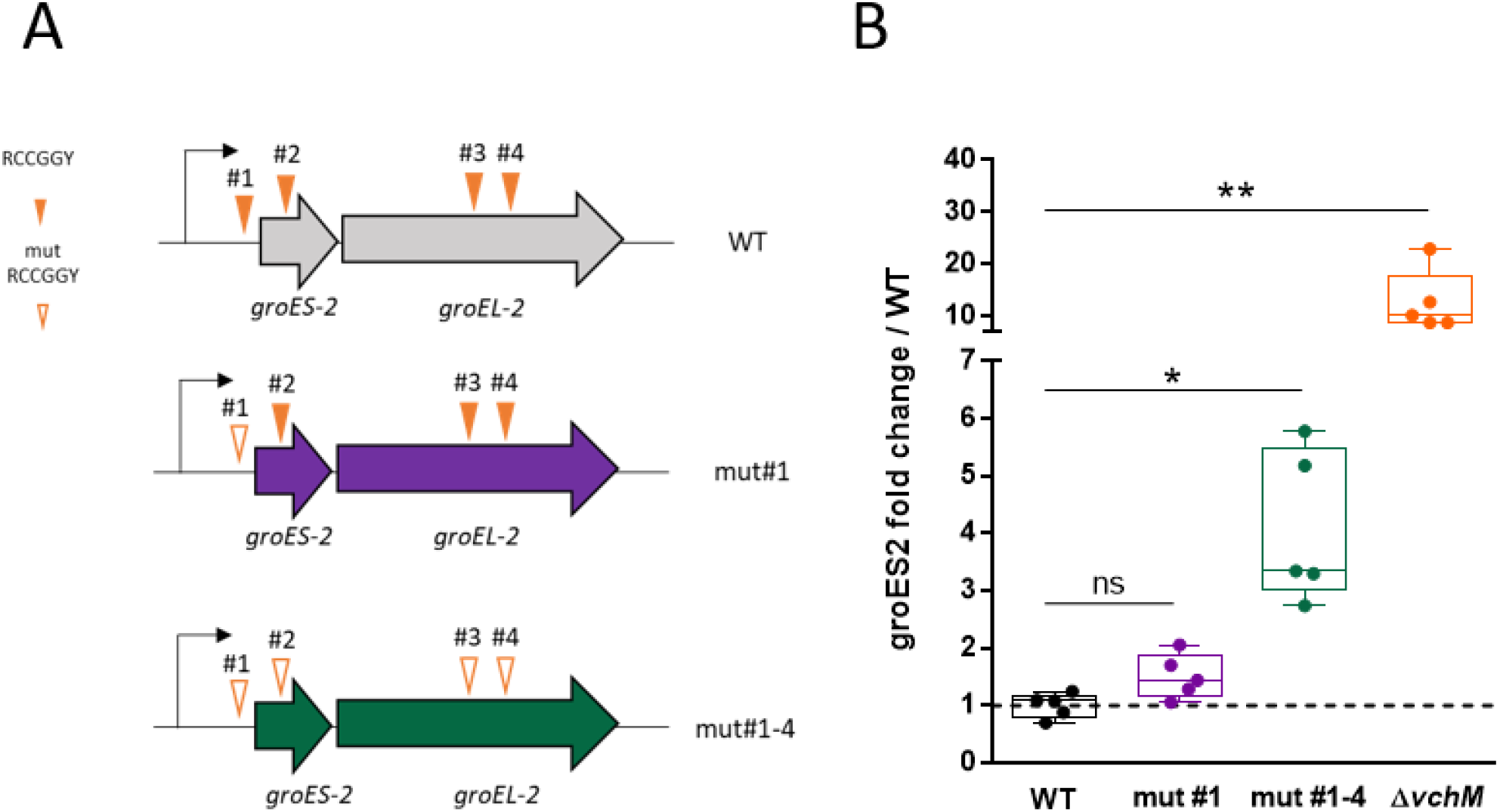
Disrupted VchM sites in *groESL-2* region leads to increased gene expression in the WT. **A.** Schematic representation of mutants with abrogated VchM sites. **B.** Relative expression of *groES-2* in the different strains at OD_600_ of 1.0. Box plots indicate the median and the 25^th^ and 75^th^ percentiles; whiskers indicate the min and max values (n= 5). Statistical significance was determined by Kruskal-Wallis test with Dunn’s post-hoc test for multiple comparisons. * P<0.05, ** P<0.01, ns = not significant

## DISCUSSION

Antimicrobial resistance (AMR) is currently one of the biggest threats to global health (35). It is thus urgent not only to find new and alternative ways to fight bacterial infections but also to understand how bacteria adapt to the presence of antibiotics and study the molecular mechanisms they use to circumvent antibiotic action.

In this study, we establish a previously unknown link between VchM-mediated DNA methylation and aminoglycoside susceptibility in the human pathogen *V. cholerae*. VchM is a relatively understudied orphan DNA methyltransferase only found in *V. cholerae* species, known to methylate the first cytosine at 5′-RCCGGY-3′ DNA motifs (16, 23). VchM is necessary for the optimal growth of *V. cholerae*, both *in vitro* and *in vivo*, and it was shown to repress the expression of a gene important for cell envelope stability through direct DNA methylation (23).

Here we show that despite the growth defect in stress-free medium, cells lacking VchM are also less susceptible to aminoglycoside toxicity. Specifically, we show that these cells have a higher relative fitness in presence of low AG concentrations. The reason for this can be inferred from the growth curves in presence of subMIC TOB (Fig 1A) where it is clear that small increments in TOB concentration lead to a higher toxicity in the WT strain when compared to the Δ*vchM*. Moreover, even though the MIC values for the tested AGs are the same in both strains, Δ*vchM* displays a higher tolerance to lethal aminoglycoside treatment (Fig 2).

Aminoglycosides are a well-known class of antimicrobial drugs that cause disruption of the translation process and consequently protein misfolding (14, 36). The exact mechanism underlying the bactericidal activity of aminoglycosides has been subject of debate in the literature (37) but it is generally accepted that killing by AGs involves i) the uptake of the AG into the cytoplasm (11, 38, 39) and ii) membrane disruption mediated by insertion of misfolded proteins in the membrane as consequence of AG binding to the ribosomes and disruption of translational fidelity (11–13, 40). Indeed, mechanisms modulating aminoglycoside tolerance/resistance in different bacterial species (in exponential or stationary phase) have been shown to be associated either to AG uptake (41–44) or to translational fidelity and proteostasis (14, 29, 40, 45). Our results revealed a higher relative abundance of *groESL-2* transcripts in bacterial cells lacking VchM, which led us to hypothesize that such increased expression of these chaperonins could underlie the high tolerance to AGs observed in this mutant, as it had been previously observed in *E. coli* (29). In fact, we show that stationary phase cells lacking both *vchM* and *groESL-2* genes have similar or even lower tolerance to lethal AG treatment compared to the WT strain. However, we could not observe a significant increase in tolerance upon overexpression of *groESL-2* in the WT strain (S5 Fig), suggesting that high *groESL-2* levels alone do not explain the high tolerance to lethal AG treatment. Instead, it is possible that high levels of GroESL-2 chaperone system counteract AG-mediated misfolding of specific substrates present only in cells devoid of VchM.

*V. cholerae* harbors two copies of *groESL* operon in its genome, thus belonging to the group of 30% of bacterial species that contains multiple copies of these chaperonins (46, 47). An interesting question to ask is whether these extra copies of chaperonins are functionally redundant or have a more specialized role in the cell, as it had been observed for *Myxococcus xanthus* (46–48). Supporting the latter hypothesis, we show here that the high tolerance observed in Δ*vchM* is dependent on the high expression of *groESL-2* but not on the high expression of *groESL-1* (Fig 4B). Amino acid identity comparison between these proteins suggests that *Vc* GroESL-1 is likely the orthologue of the housekeeping GroESL of *E. coli* whereas *Vc* GroESL-2, thought to have appeared by duplication in *V. cholerae* (30), differ equally from both (S2 Fig). Thus, we speculate that *V. cholerae* GroESL-2 constitutes an alternative chaperone system capable of helping the folding of specific substrates important for survival to specific stresses. Interestingly, even though essential for the higher tolerance to lethal aminoglycoside treatment, *groESL-2* is not involved in the increased relative fitness of the Δ*vchM* mutant in presence of subMIC doses of aminoglycosides (Fig 4C). This suggests that the mechanisms operating in Δ*vchM* cells that increase their relative fitness during growth in subMIC AGs are not the same that increase their tolerance to lethal doses of these drugs. In fact, it has been recently shown that the type of translation errors occurring at lower streptomycin (another AG) concentrations differ from those found in high concentrations of this aminoglycoside, with the latter being associated to a higher misfolding propensity (12). Thus, it seems plausible that, in Δ*vchM* cells, the higher expression of *groESL-2* is likely to be more important at high concentrations of AGs, when the abundance of misfolded proteins tend to increase. The mechanisms driving Δ*vchM* higher relative fitness at lower doses of aminoglycosides remain to be elucidated in future work.

DNA methylation controls gene expression through modulation of protein-DNA interactions (49). In most of the cases, the methylated base interferes with the binding of transcription factors and/or the RNA polymerase at the regulatory region of a gene, affecting transcription (50–52). However, there is also evidence that the presence of methylated DNA bases that occur along the coding region of genes could also directly affect their expression in bacteria, even though the precise mechanism is still unknown (20, 22, 23). In eukaryotes, cytosine methylation tends to repress gene expression. A recent study shedding light on how cytosine methylation affect DNA mechanical properties shows that cytosine methylation stabilizes the DNA helix and slows transcription in eukaryotic cells (53). Thus, a similar m5C-mediated transcriptional hindrance is likely to happen also in prokaryotes. Here we support this view by showing that abrogation of VchM-dependent methylation of cytosines at the four RCCGGY motifs in *groESL-2* region increased its expression in WT cells (Fig 5B). However, this in unlikely the sole mechanism responsible for the high expression of *groESL-2* genes in Δ*vchM* cells, as this mutant has even higher expression levels of *groESL-2*. It is possible that the pleiotropic effects resulting from VchM deficiency also affect, indirectly, the expression of these genes through regulation of a specific transcription factor.

Our work shows that a *V. cholerae* deletion mutant of the orphan DNA methyltransferase VchM have a general higher tolerance towards aminoglycosides. It remains to be explored whether *V. cholerae* WT cells can modulate VchM expression and, consequently, alter the levels of cytosine methylation. Bisulfite sequencing analysis of *V. cholerae* genome shows that all cytosines within RCCGGY motifs were methylated in *V. cholerae*, during exponential and stationary phases, with the exception of three of these sites which had been previously shown to be constantly undermethylated in this species (54). However, these studies were conducted in cells cultured in LB stress-free media or collected from frozen rabbit cecal fluid, and thus may not reflect the m5C profile of *V. cholerae* during other stress conditions. Moreover, bisulfite sequencing allows for cytosine methylation analysis of the total population at a specific time and thus it is not suitable to detect potential transient changes in small subpopulations of cells. Such changes could be mediated, for example, by altering the levels of VchM through gene expression. Little is known about *vchM* regulation but it was recently shown that the *V. cholerae* quorum sensing low density transcriptional regulator AphA is able to bind the *vchM* region (55) leaving the possibility that *vchM* may be regulated by quorum sensing. Moreover, *vchM* was previously found to be differentially expressed between different stages of human infection (56), suggesting the possibility that modulation of cytosine methylation levels can be adaptative during *V. cholerae*’s life cycle. In line with our work, lowering VchM levels could lead to a trade-off, where low m5C levels would be detrimental for fitness in stress-free contexts, but highly advantageous in presence of specific stress conditions, such as antibiotic exposure.

## MATERIALS AND METHODS

### Strains, media and culture conditions

*V. cholerae* was routinely cultured at 37°C in Mueller-Hinton (MH) medium. Plasmids were introduced in *V. cholerae* by electrotransformation. Strains containing the pSC101 plasmid were grown in presence of 100 μg/mL carbenicilin for plasmid maintenance. All *V. cholerae* mutant strains are derived from *Vibrio cholerae* serotype O1 biotype El Tor strain N16961 hapR+. Mutants were constructed by homologous recombination after natural transformation or with a conjugative suicide plasmid as previously described (15, 57–59). Primers, strains and plasmids used in this study, and their constructions, are listed in Table S2. For routine cloning we used chemically competent *E. coli* One Shot^®^ TOP10 (Invitrogen). All strains and plasmids were confirmed by sanger sequencing.

### Mutation of RCCGGY sites #1-4 in *groESL-2* region

In order to mutate all four RCCGGY sites present in *groESL-2* (*vca0819-0820*) region we generated a DNA fragment (S2 Table) with these sites containing the following nucleotide changes: #1-ACCGGC changed to A**<u>T</u>**CGGC; #2-ACCGGC changed to AC**<u>G</u>**GGC; #3-GCCGGC changed to GC**<u>G</u>**GGC and #4-ACCGGC changed to AC**<u>G</u>**GGC. This fragment was then introduced in *V. cholerae* at the endogenous locus by allelic replacement as described in Table S2.

### Growth curves

Overnight cultures from single colonies were diluted 1:100 in Mueller-Hinton (MH) rich media or MH + subMIC antibiotics at different concentrations, in 96-well microplates. OD_600_ was measured in a Tecan Infinite plate reader at 37°C, with agitation for 20 hours. Measurements were taken every 10 minutes.

### MIC determination

MICs were determined by microtiter broth dilution method (60) with an initial inoculum size of 10^5^ CFUs/mL. The MIC was interpreted as the lowest antibiotic concentration preventing visible growth.

### Neo-Cy5 uptake

Quantification of fluorescent neomycin (Neo-cy5) uptake was performed as described (61). Neo-cy5 is an aminoglycoside coupled to the fluorophore Cy5, and has been shown to be active against Gram-bacteria (27). Briefly, overnight cultures were diluted 100-fold in rich MOPS (Teknova EZ rich defined medium). When the bacterial cultures reached an OD_600_ of 0.25, they were incubated with 0.4 μM of Cy5 labeled Neomycin for 15 minutes at 37°C. 10 μL of the incubated culture were then used for flow cytometry, diluting them in 250 μL of PBS before reading fluorescence. Flow cytometry experiments were performed as described (62). For each experiment, 100000 events were counted on the Miltenyi MACSquant device.

### Competitions experiments

Overnight cultures from single colonies of lacZ^-^ and lacZ^+^ strains were washed in PBS (Phosphate Buffer Saline) and mixed 1:1 (500μl + 500μl). At this point 100μl of the mix were serial diluted and plated in MH agar supplemented with X-gal (5-bromo-4-chloro-3-indolyl-β-D-galactopyranoside) at 40 μg/mL to assess T0 initial 1:1 ratio. At the same time, 10 μl from the mix were added to 2 mL of MH or MH supplemented with subMIC tobramycin at 0.6μg/mL and incubated with agitation at 37°C for 20 hours. Cultures were then diluted and plated in MH agar plates supplemented with X-gal. Plates were incubated overnight at 37°C and the number of blue and white CFUs was assessed. Competitive index was calculated by dividing the number of blue CFUs (lacZ^+^ strain) by the number of white CFUs (lacZ^-^ strain).

### Survival assays

Bacterial cultures from single colonies were cultured at 37°C for 16 h with agitation in 10 mL of MH medium. Aliquots from these cultures were removed, serial diluted and plated in MH agar plates to assess CFUs formation prior antibiotic treatment (T0). In addition, 5 mL of these aliquots were subjected to antibiotic treatment and incubated with agitation at 37°C. At the indicated time points, 500uL of these cultures were collected, washed in PBS, serial diluted and plated in MH agar plates. The plates were then incubated overnight at 37°C. Survival at each time point was determined by dividing the number of CFUs/mL at that time point by the number of CFUs/mL prior treatment. Antibiotics were used at the following final concentrations: 20 μg/mL Tobramycin (TOB) and 10 μg/mL Gentamicin (GEN). Experiments were repeated at least two to three times.

### Digital qRT-PCR

For RNA extraction, overnight cultures of three biological replicates of strains of interest were diluted 1:1000 in MH media and grown with agitation at 37°C until an OD_600_ of 0.3 (exponential phase) or an OD_600_ of 1.0 or 2.0 (stationary phase). 0.5 mL of these cultures were centrifuged and supernatant removed. Pellets were homogenized by resuspension with 1.5 mL of cold TRIzol™ Reagent. Next, 300 μL chloroform were added to the samples following mix by vortexing. Samples were then centrifuged at 4°C for 10 minutes. Upper (aqueous) phase was transferred to a new 2mL tube and mixed with 1 volume of 70% ethanol. From this point, the homogenate was loaded into a RNeasy Mini kit (Quiagen) column and RNA purification proceeded according to the manufacturer’s instructions. Samples were then subjected to DNase treatment using TURBO DNA-free Kit (Ambion) according to the manufacturer’s instructions. RNA concentration of the samples was measured with NanoDrop™ spectrophotometer and diluted to a final concentration of 1-10 ng/μL.

qRT-PCR reactions were prepared with 1 μL of diluted RNA samples using the qScript™ XLT 1-Step RT-qPCR ToughMix (Quanta Biosciences, Gaithersburg, MD, USA) within Sapphire chips. Digital PCR was conducted on a Naica Geode (programmed to perform the sample partitioning step into droplets, followed by the thermal cycling program suggested in the user’s manual. Primer and probe sequences used in digital qRT-PCR reaction are listed in Table S3. Image acquisition was performed using the Naica Prism3 reader. Images were then analyzed using Crystal Reader software (total droplet enumeration and droplet quality control) and the Crystal Miner software (extracted fluorescence values for each droplet). Values were normalized against expression of the housekeeping gene *gyrA* as previously described (31).

### RNA-seq

For RNA extraction, overnight cultures of three biological replicates of WT and Δ*vchM* strains were diluted 1:100 in MH medium and grown with agitation at 37°C until cultures reach an OD_600_ of 2.0. Total RNA extraction, library preparation, sequencing and analysis were performed as previously described (63). The data for this RNA-seq study has been submitted in the GenBank Sequence Read Archive (SRA) under project number PRJNA509113.

## ACKNOWLEDGMENTS

We are thankful to Manon Lang for her valuable help with the neo-cy5 uptake experiments and Sebastian Aguilar Pierlé for all the help with RNAseq analysis. We also thank Evelyne Krin for help with molecular cloning procedures and João Gama for helpful comments on the manuscript.

## SUPPORTING INFORMATION

**S1 Table. Differentially regulated genes in ΔvchM strain**

**S2 Table. Strains, plasmids and primers used in this study**

**S3 Table. Primer and probe sequences used in digital qRT-PCR**

**S1 Fig. Neo-cy5 uptake is not increased in Δ*vchM*.** Percentage of neo-cy5 positive cells analyzed by flow cytometry after incubation with fluorescent marked neomycin. Means and SD are represented, n=3.

**S2 Fig. Comparison of GroESL proteins from *E. coli* and *V. cholerae*.** Amino acid identity between GroES and GroEL proteins of *E. coli* MG1655 (*Eco*) and *V. cholerae* O1 El Tor N16961 (*Vch*) computed by BLASTP. Values represent percentage identity between proteins.

**S3 Fig. Deletion of *groESL-2* does not increase susceptibility to tobramycin. A.** *In vitro* competitions of WT and Δ*groESL-2* strains against isogenic Δ*lacZ* reference strain in absence or presence of tobramycin (TOB) at 0.6 μg/ml; n=3, error bars indicate SD. **B.** Survival of stationary-phase WT and Δ*groESL-2* cells exposed to 20X MIC of tobramycin. n=3, error bars indicate SD.

**S4 Fig. Mutation of VchM sites in *groESL-2* region fails to affect gene expression of the operon in absence of VchM.** Relative expression of *groES-2 and groEL*-2 genes in the indicated strains grown at OD_600_ 1.0. n=3, error bars indicate SD.

**S5 Fig. Overexpression of *groESL-2* in the WT does not increase tolerance to tobramycin.** Survival (after 7 hours TOB treatment) of WT strain carrying a control plasmid or a plasmid overexpressing *groESL-2* genes. n=6, error bars indicate SD.

