## Supplementary material for "Deficiency in cytosine DNA methylation leads to high chaperonin expression and tolerance to aminoglycosides in *Vibrio cholerae*": S1 Table.pdf

**Table S1. Differentially regulated genes in *ΔvchM* strain**

| Locus | Name | Annotation | Fold change *ΔvchM*/WT | p-value |
|---|---|---|---|---|
| **UPREGULATED** | | | | |
| ***vca1028*** | *lamB* | maltoporin | 52,81 | 0,0E+00 |
| ***vca0707*** | *uhpC* | MFS transporter | 8,76 | 8,6E-06 |
| ***vc1368*** | | conserved hypothetical protein | 8,18 | 0,0E+00 |
| ***vca0819*** | ***groES-2*** | **chaperonin** | **6,78** | **0,0E+00** |
| ***vca0200*** | | putative ATPase | 6,10 | 0,0E+00 |
| ***vca0957*** | *glcB* | putative Malate synthase | 5,97 | 0,0E+00 |
| ***vca0199*** | | conserved hypothetical protein | 5,32 | 5,2E-09 |
| ***vca1027*** | *malM* | putative Maltose operon periplasmic protein MalM | 4,14 | 0,0E+00 |
| ***vca0201*** | | conserved hypothetical protein | 3,97 | 3,4E-11 |
| ***vca0946*** | *malK* | Maltose/maltodextrin import ATP-binding protein malK | 3,91 | 0,0E+00 |
| ***vca0469*** | *higA2* | plasmid stabilization system HigA protein (antitoxin of TA system HigB-HigA) | 3,91 | 0,0E+00 |
| ***vca0468*** | *higB2* | plasmid stabilization system HigB protein (toxin of TA system HigB-HigA) | 3,90 | 2,3E-10 |
| ***vca0945*** | *malE* | maltose ABC transporter periplasmic binding protein | 3,90 | 0,0E+00 |
| ***vc1819*** | *aldA* | Aldehyde dehydrogenase | 3,84 | 1,7E-12 |
| ***vc1204*** | *hutG* | putative Formimidoylglutamase HutG (histidine utilization) | 3,80 | 2,2E-16 |
| ***vc1203*** | *hutU* | Urocanate hydratase (histidine utilization) | 3,60 | 0,0E+00 |
| ***vca0820*** | ***groEL-2*** | **chaperonin** | **3,54** | **2,2E-16** |
| ***vca0692*** | *secF* | fragment of putative SecD/SecF/SecDF export membrane protein (part 2) | 3,48 | 3,2E-08 |
| ***vc1809*** | *alpA* | putative Prophage CP4-57 regulatory protein (AlpA) | 3,37 | 1,8E-09 |
| ***vc1205*** | *hutI* | Imidazolonepropionase (histidine utilization) | 3,24 | 1,1E-11 |
| ***vc1740*** | *fadE* | acyl coenzyme A dehydrogenase | 3,22 | 5,6E-15 |
| ***vca0944*** | *malF* | maltose ABC transporter membrane subunit MalF | 3,18 | 0,0E+00 |
| ***vc1142*** | *cspD* | Cold shock-like protein cspD | 2,88 | 0,0E+00 |
| ***vc2530*** | *hpf* | putative ribosome hibernation promoting factor HPF/sigma 54 modulation protein | 2,83 | 0,0E+00 |
| ***vca0948*** | *yaeO* | putative Rho-specific inhibitor of transcription termination (YaeO) | 2,78 | 0,0E+00 |
| ***vca0998*** | *xenB* | Xenobiotic reductase B | 2,73 | 4,4E-06 |

| | | | | |
|---|---|---|---|---|
| ***vc0486*** | *srlR* | putative Transcriptional regulator of sugar metabolism; DeoR family transcriptional regulator | 2,65 | 9,9E-14 |
| ***vc1874*** | | putative SpoVR family protein | 2,64 | 0,0E+00 |
| ***vc1202*** | *hutH* | Histidine ammonia-lyase | 2,61 | 0,0E+00 |
| ***vc2265*** | *groES-1* | chaperonin | 2,60 | 0,0E+00 |
| ***vc2615*** | | conserved hypothetical protein | 2,55 | 4,6E-10 |
| ***vc1678*** | *pspA* | Phage shock protein A | 2,48 | 4,6E-10 |
| ***vc1183*** | | putative 2OG-Fe dioxygenase | 2,43 | 0,0E+00 |
| ***vca0470*** | | putative Acetyltransferase | 2,43 | 2,1E-08 |
| ***vca0324*** | *relB1* | Plasmid stabilization system protein RelB (Anti Toxin of TA system RelB-RelE) | 2,43 | 1,1E-11 |
| ***vca0886*** | *kbl* | glycine C-acetyltransferase | 2,40 | 2,3E-08 |
| ***vca0185*** | *arfA* | putative Stalled ribosome alternative rescue factor ArfA | 2,39 | 3,0E-13 |
| ***vca0958*** | | putative transcriptional regulator | 2,38 | 3,9E-10 |
| ***vca0013*** | *malP* | maltodextrin phosphorylase | 2,37 | 3,9E-11 |
| ***vca0881*** | | conserved hypothetical protein | 2,36 | 1,1E-13 |
| ***vca0551*** | | conserved hypothetical protein | 2,35 | 0,0E+00 |
| ***vca0391*** | *higB1* | Plasmid stabilization system HigB protein (Toxin of TA system HigB-HigA) | 2,34 | 1,5E-10 |
| ***vca0923*** | *mlp37* | chemoreceptor Mlp37 | 2,34 | 2,2E-12 |
| ***vc2361*** | *grcA* | Autonomous glycyl radical cofactor | 2,30 | 0,0E+00 |
| ***vca0392*** | *higA1* | Plasmid stabilization system HigA protein ( Antitoxin of TA system HigB-HigA) | 2,28 | 0,0E+00 |
| *vc0665* | *vpsR* | Fis family transcriptional regulator (sigma-54 dependent transcriptional regulator) | 2,28 | 0,0E+00 |
| ***vca0720*** | *hnoX* | Heme-Nitric Oxide/Oxygen Binding Protein (H-NOX) | 2,27 | 5,1E-10 |
| ***vc1433*** | *uspE* | universal stress protein UspE | 2,26 | 0,0E+00 |
| ***vc1248*** | *tar* | putative Methyl-accepting chemotaxis protein | 2,25 | 0,0E+00 |
| ***vc2264*** | *groEL-1* | chaperonin | 2,24 | 0,0E+00 |
| ***vc0734*** | *aceB* | Malate synthase A | 2,24 | 1,4E-06 |
| ***vc2507*** | | PhoH family protein | 2,23 | 0,0E+00 |
| ***vc1344*** | *hppD* | 4-hydroxyphenylpyruvate dioxygenase | 2,22 | 6,9E-15 |
| ***vca0219*** | *hlyA* | Hemolysin | 2,20 | 5,7E-14 |
| ***vca0159*** | | putative Universal stress protein family 1 | 2,17 | 2,9E-15 |
| ***vca0987*** | *ppsA* | phosphoenolpyruvate synthase | 2,15 | 2,6E-09 |

| | | | | |
|---|---|---|---|---|
| ***vca0359*** | *parE2* | Plasmid stabilization system ParE protein (toxin of TA system ParD-ParE) | 2,14 | 9,4E-10 |
| ***vca0004*** | | conserved hypothetical protein | 2,14 | 0,0E+00 |
| ***vc1222*** | *ihfA* | integration host factor subunit α | 2,13 | 0,0E+00 |
| ***vc2758*** | *fadB* | fatty acid oxidation complex subunit alpha FadB | 2,10 | 6,1E-10 |
| ***vc2704*** | | conserved hypothetical protein | 2,10 | 8,9E-09 |
| ***vc1962*** | *nlpE* | copper resistance protein NlpE N-terminal domain-containing protein | 2,08 | 3,9E-09 |
| ***vca0278*** | *glyA* | serine hydroxymethyltransferase | 2,04 | 0,0E+00 |
| ***vca0280*** | *gcvT* | glycine cleavage system aminomethyltransferase | 2,04 | 4,4E-16 |
| ***vc0976*** | *qmcA* | putative Membrane protease subunits, stomatin/prohibitin homolog qmcA | 2,03 | 6,5E-06 |
| ***vc1539a*** | | hypothetical protein | 2,02 | 6,9E-10 |
| ***vc0666*** | | conserved hypothetical protein | 2,02 | 1,8E-09 |
| ***vc2144*** | *flaF* | Polar flagellin F | 2,01 | 3,0E-13 |
| **DOWNREGULATED** | | | | |
| **Locus** | **Name** | **Annotation** | **Fold change Δ*vchM*/WT** | **p-value** |
| ***vca0198*** | *vchM* | Cytosine-specific DNA methyltransferase | -12,22 | 1,4E-24 |
| ***vca0933*** | *cspE* | transcription antiterminator and regulator of RNA stability | -8,41 | 1,0E-07 |
| ***vca0017*** | *hcp* | Haemolysin co-regulated protein (putative Type VI secretion system effector, Hcp1) | -3,20 | 1,3E-10 |
| ***vc2383*** | *ilvY* | putative Transcriptional regulator, LysR family | -3,14 | 5,9E-27 |
| ***vca0804*** | *deaD* | Cold-shock DEAD box protein A | -3,12 | 6,5E-49 |
| ***vca0874*** | | conserved hypothetical protein | -3,10 | 1,1E-05 |
| ***vc1704*** | *metE* | 5-methyltetrahydropteroyltriglutamate--homocysteine methyltransferase | -3,02 | 7,6E-21 |
| ***vc0768*** | *guaA* | GMP synthetase | -2,91 | 2,7E-72 |
| ***vca0680*** | *napC* | Cytochrome c-type protein napC | -2,87 | 2,4E-07 |
| ***vca0679*** | *napB* | Diheme cytochrome c napB | -2,82 | 2,1E-06 |
| ***vca0935*** | | hypothetical protein | -2,76 | 1,2E-10 |
| ***vc1261*** | | putative Sugar efflux transporter B | -2,63 | 3,9E-07 |
| ***vc0069*** | *mdtL* | putative FLORFENICOL EXPORTER | -2,53 | 8,1E-14 |
| ***vca1002*** | | AzlC family ABC transporter permease | -2,49 | 2,0E-08 |
| ***vc1415*** | *hcp* | type VI secretion system secreted protein Hcp | -2,48 | 1,0E-06 |
| ***vc0433*** | *arcD* | Arginine/ornithine antiporter | -2,48 | 1,9E-06 |

| | | | | |
|---|---|---|---|---|
| ***vc1393*** | *sugE* | Quaternary ammonium compound-resistance protein sugE | -2,43 | 3,4E-22 |
| ***vc1035*** | | conserved hypothetical protein | -2,40 | 6,3E-06 |
| ***vc1608*** | | ABC transporter permease | -2,38 | 2,8E-11 |
| ***vca0250*** | | Alpha-amylase | -2,37 | 7,2E-08 |
| ***vc0767*** | *guaB* | inositol-5-monophosphate dehydrogenase | -2,31 | 3,0E-22 |
| ***vc1239*** | *cobU* | cobinamide-P guanylyltransferase / cobinamide kinase | -2,28 | 1,2E-09 |
| ***vc0290*** | *fis* | DNA-binding protein fis | -2,27 | 1,4E-04 |
| ***vc2227*** | *purN* | phosphoribosylglycinamide formyltransferase 1 | -2,25 | 2,6E-10 |
| ***vc2384*** | | TSUP family transporter | -2,24 | 8,1E-16 |
| ***vca1003*** | | AzlD domain-containing protein | -2,23 | 1,5E-11 |
| ***vca0678*** | *napA* | Periplasmic nitrate reductase | -2,22 | 2,6E-10 |
| ***vca1031*** | | Methyl-accepting chemotaxis protein | -2,21 | 7,0E-11 |
| ***vc1843*** | *cydB* | Cytochrome d ubiquinol oxidase subunit 2 | -2,21 | 5,9E-13 |
| ***vc0191*** | *rhtC* | putative Lysine/Homoserine/Threonine exporter protein | -2,20 | 2,6E-10 |
| ***vc1609*** | | ABC transporter permease | -2,19 | 2,2E-10 |
| ***vc0005*** | *yidD* | membrane protein insertion efficiency factor | -2,16 | 4,9E-10 |
| ***vc0291*** | *dusB* | tRNA-dihydrouridine synthase B | -2,16 | 3,8E-16 |
| ***vc1855*** | *dinG* | ATP-dependent DNA helicase DinG | -2,15 | 7,7E-07 |
| ***vc2382*** | | conserved hypothetical protein | -2,15 | 3,1E-07 |
| ***vc1842*** | *cydX* | cytochrome bd-I oxidase subunit CydX | -2,14 | 1,8E-06 |
| ***vc1629*** | | ABC transporter permease | -2,13 | 3,0E-06 |
| ***vc0706*** | *raiA* | Ribosome-associated inhibitor A | -2,13 | 1,1E-08 |
| ***vc2047*** | | SDR family oxidoreductase | -2,11 | 3,4E-23 |
| ***vc1174*** | *trpE* | Anthranilate synthase component 1 | -2,09 | 9,9E-18 |
| ***vc1240*** | | histidine phosphatase family protein | -2,09 | 1,5E-06 |
| ***vc0940*** | | conserved hypothetical protein | -2,07 | 1,3E-07 |
| ***vc1487*** | | glutaredoxin family protein | -2,06 | 8,0E-06 |
| ***vca0214*** | *emrD* | multidrug efflux pump EmrD | -2,05 | 6,0E-12 |
| ***vca0269*** | | aspartate aminotransferase family protein | -2,04 | 2,2E-13 |
| ***vc2647*** | *aphA* | Transcriptional regulator PadR family | -2,04 | 9,6E-56 |
| ***vc2226*** | *purM* | Phosphoribosylformylglycinamidine cyclo-ligase | -2,02 | 9,3E-10 |
| ***vc0651*** | | putative Peptidase U32 | -2,02 | 2,4E-13 |
| ***vc1630*** | | putative SalX, ABC-type antimicrobial peptide transport system, ATPase component | -2,02 | 2,5E-19 |
| ***vca0684*** | | MFS transporter | -2,02 | 9,3E-14 |

| | | | | |
|---|---|---|---|---|
| ***vc0717*** | *yegQ* | tRNA 5-hydroxyuridine modification protein YegQ | -2,01 | 4,2E-09 |
| ***vc1856*** | | TSUP family transporter | -2,00 | 7,2E-10 |
| ***vc0987*** | *hemH* | ferrochelatase | -2,00 | 1,9E-17 |
