## Supplementary material for "Deficiency in cytosine DNA methylation leads to high chaperonin expression and tolerance to aminoglycosides in *Vibrio cholerae*": S2 Table.pdf

### Table S2. Strains, plasmids and primers used in this study

| Strain | Lab Strain number | Source/Construction |
|---|---|---|
| ***Vibrio cholerae*** | | |
| N16961 hapR+ WT strain | F606 | Gift from Melanie Blokesch |
| Δ*vchM* (vca0198) | H507 | PCR amplification of 500bp up and down genomic regions of VCA0198 using primers ZIP136/137 and ZIP138/139. PCR amplification of aadA7 conferring spectinomycin resistance on pAM34 using ZB47/48. PCR assembly of the VCA0198::spec fragment using ZIP136/139 and allelic exchange by natural transformation in F606. |
| Δ*lacZ* | K329 | Lab collection |
| Δ*groESL-2* (*vca0819-0820*) | M958 | allelic exchange by integration and excision of conjugative suicide plasmid pMP7 (pM340) replacing the gene with frt::kan::frt |
| Δ*vchM* Δ*groESL-2* | N330 | PCR amplification of 500bp up and down genomic regions of VCA0198 using primers ZIP136/137 and ZIP138/139. PCR amplification of aadA7 conferring spectinomycin resistance on pAM34 using ZB47/48. PCR assembly of the VCA0198::spec fragment using ZIP136/139 and allelic exchange by natural transformation in M958 |
| WT – VchM site #1 mutated (mut #1) | L900 | allelic exchange by integration and excision of conjugative suicide plasmid pMP7 (pL442) |
| WT – VchM sites #1-4 mutated (mut #1-4) | Q827 | allelic exchange by integration and excision of conjugative suicide plasmid pMP7 containing the *groESL-2* region with sites #1-4 mutated (pQ824) in M958 strain. |
| Δ*vchM* mut #1-4 | Q828 | allelic exchange by integration and excision of conjugative suicide plasmid pMP7 containing the *groESL-2* region with sites #1-4 mutated (pQ824) in N330 |

| Plasmids | | |
|---|---|---|
| pMP7-Δ*vca0819-0820*::kan | M340 | gibson assembly using primers MV450/451 for the amplification of pMP7 vector, primers "vca0819-8205" and "vca0819-8206" for up and down regions of the gene, and primers MV268/269 on pKD4 plasmid for the resistance gene (frt::kan::frt). |
| pMP7- C->T point mutation of VchM site #1 in 5' UTR region of vca0819-0820 | L442 | Amplification of 500bp upstream of *vca0819* with primers 5923/5922; Amplification of 500bp downstream of *vca0819* with 5924/5921; PCR assembly of the two fragments with primers 5923/5924. Note: Primers 5922 and 5921 contain a mismatch to give origin to a C->T point mutation in VchM site #1. The fragment was then cloned in a pTOPO vector and sub cloned into pMP7 using *EcoRI* restriction sites. |
| pMP7- sites #1-4 mutated in vca0819-0820 region | Q824 | A *groESL-2* fragment with mutations in VchM sites #2-4 was synthesized and cloned in a pTOPO vector. This fragment was then PCR assembled to the 5' UTR region containing site #1 mutated (from strain L900), which originated a final fragment containing the #1-4 mutated sites. This fragment was then cloned in pTOPO and sub cloned into pMP7 using *EcoRI* restriction sites. |
| pSC101-*groESL-1* | O849 | Amplification of *vc2664-2665* from *V. cholerae* gDNA with primers AFC046/AFC047. Primer AFC046 contains a Ptrc promoter. Fragment cloned in pSC101 low copy plasmid (carbenicillin resistant) using *EcoRI* restriction sites |
| pSC101-*groESL-2* | N752 | Amplification of *vca0819-0820* from *V. cholerae* gDNA with primers AFC029/AFC030. Primer AFC029 contains a Ptrc promoter. Fragment cloned in pSC101 low copy plasmid (carbenicillin resistant) using *EcoRI* restriction sites |
| pSC101-*vchM* | Q826 | Amplification of *vca0198 (vchM)* with its own promoter from *V. cholerae* gDNA with primers 5990/5911. Cloning in pTOPO vector and sub cloning in pSC101 low copy plasmid (carbenicillin resistant) using *BamHI* and *PstI* restriction sites |
| **Primers** | | **Sequence 5'-3'** |
| ZIP136 | | GCCGCCGAAGGAAAAACCGTACTATTGC |
| ZIP137 | | GCGAGCATCGTTTGTTCGCCCAGCTTCTGTATGGAACGGGTTAACTGTATCACCATACTACCTCATGG |
| ZIP138 | | CGTGAAAGGCGAGATCACCAAGGTAGTCGGCAAATAATGTCTACATGCTTCACAGCGTAGTCGC |
| ZIP139 | | TTAATTTCTCGAGTTTCAGATGC |

| | | |
|---|---|---|
| ZB47 | | CCCGTTCCATACAGAAGCTGGGCGAACAAACGATGCTCGC |
| ZB48 | | GACATTATTTGCCGACTACCTTGGTGATCTCGCCTTTCACG |
| vca0819-8205 | | CTATTATTTAAACTCTTTCCGTTTTGCCTT |
| vca0819-8206 | | TACGTAGAATGTATCAGACTCGCCCAAGGA |
| 5921 | | AAACAATCCTA**T**CGGCCTTTTATC |
| 5922 | | GATAAAAGGCCG**A**TAGGATTGTTT |
| 5923 | | ACTTTGATGGTACGCGCGATG |
| 5924 | | GATTTATTGAGCACAACATGGCG |
| AFC029 | | GTAAGTGAATTCTTGACAATTAATCATCCGGCTCGTATAATGTGTGGAATTGTGAGCGGATAACAATTTCACACAGGAAACAGCGCCGCATGAATATTCGTCCTTTACATG |
| AFC030 | | GTAAGTGAATTCATTACGCCGCAGACTCTTTGTC |
| AFC046 | | GTAAGTGAATTCTTGACAATTAATCATCCGGCTCGTATAATGTGTGGAATTGTGAGCGGATAACAATTTCACACAGGAAACAGCGCCGCATGAATATTCGTCCATTACATGAC |
| AFC047 | | GTAAGTGAATTCCTGCTAAGGGGGATGATTACA |
| 5990 | | GTTTTTGCTGCCGTCTGCTA |
| 5911 | | GTAGTCGACCCTTTTTACAAACTTTCTAGA |
