## Supplementary material for "Deficiency in cytosine DNA methylation leads to high chaperonin expression and tolerance to aminoglycosides in *Vibrio cholerae*": S3 Table.pdf

### Table S3. Primer and probe sequences used in digital qRT-PCR

| Target | Primers (5'-3') | Probes (5'-3') |
|---|---|---|
| *gyrA* | AATGTGCTGGGCAACGACTG<br>GAGCCAAAGTTACCTTGGCC | [Cy5]-CACCCTCATGGTGACAGTGCGGTTT-[BHQ2] |
| *groES-1* | CGTAGCTTTCTGCGAAGATC<br>TGGTGGAATTGTTCTAACTG | [HEX] -AGCTCCAAAGGTTGAACTGAACCGTTCTCTA-[BHQ1] |
| *groES-2* | GGCGACCAGATCATTTTCAAC<br>TCTACAATCGCTAACACATCAG | [FAM] - TGGACGGTAAAGAGTATCTGATCCTCTCC-[BHQ1] |
| *groEL-2* | TGCTATCGTTCAGAATCAACC<br>TTTACTTCTCGCTCCCAATCG | [HEX] –AAGCTTACCAGCATAGAACTTGCCAGAGAT-[BHQ1] |
