## Supplementary figures and images for "Deficiency in cytosine DNA methylation leads to high chaperonin expression and tolerance to aminoglycosides in *Vibrio cholerae*"

### S1 Fig.tif

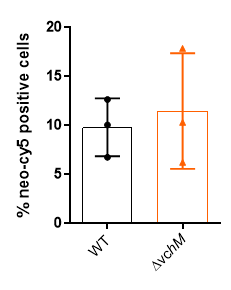

### S2 Fig.tif

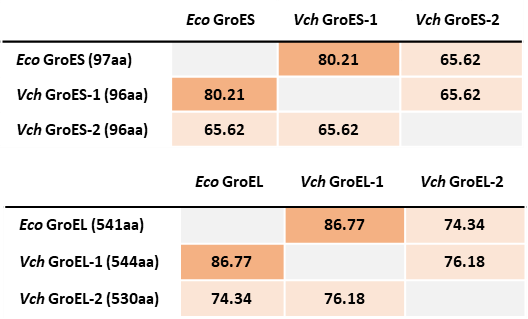

### S3 Fig.tif

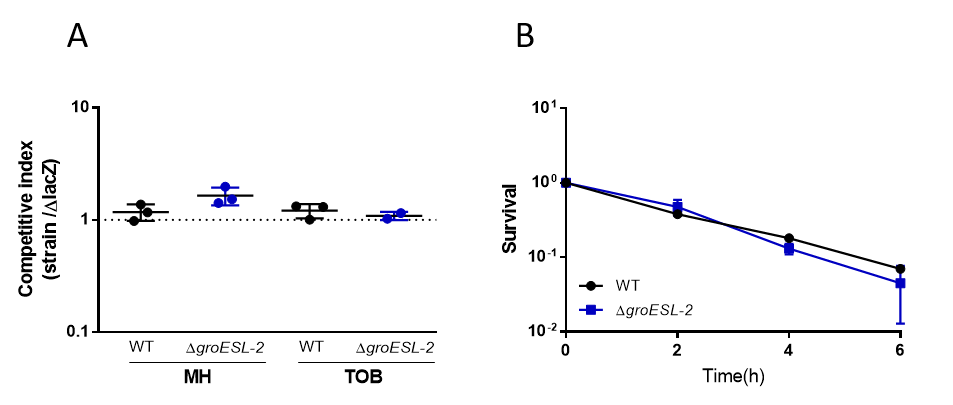

### S4 Fig.tif

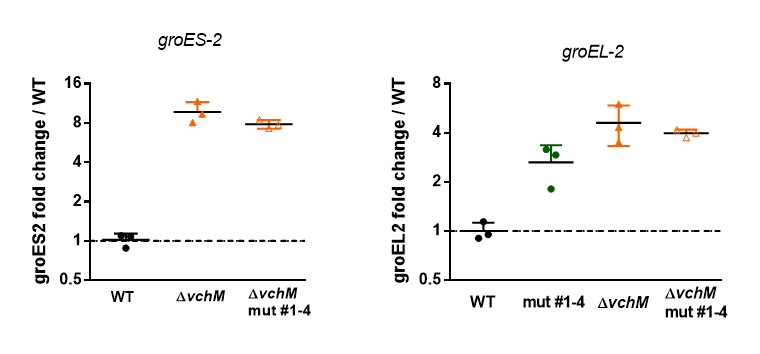

### S5 Fig.tif

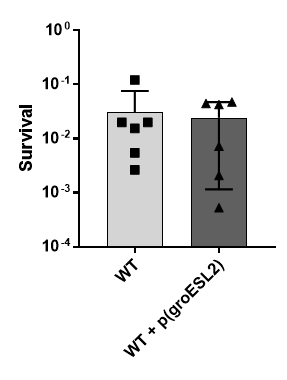
